# Transgenic mouse facial nerve model of synkinesis: Original Research

**DOI:** 10.1101/2019.12.27.877795

**Authors:** Mostafa M Ahmed, Alexander Deich, Grace Balfour, Adrienne Laury, Arkady Lorin, David Silliman, Richard L. Williams

## Abstract

**Hypothesis:** Our central hypothesis is that inhibition of Schwann cell de-differentiation, in the post-injury setting, will reduce synkinesis and improve facial muscle function

**Background:** No therapies exist to improve the accuracy of facial nerve regeneration. Following peripheral nerve injury, adult reactive Schwann cells de-differentiate and express glial fibrillary acidic protein (GFAP); suggesting that reactive Schwann cells impact axonal regeneration precision.

**Methods:** Transgenic GFAP-thymidine kinase (TK) mouse model was employed, allowing selective downregulation of reactive GFAP expressing Schwann cells on exposure of 7 day osmotic pump delivery of ganciclovir. Adult female transgenic GFAP-TK mice had right facial nerve transected and then immediately repaired with tissue glue, they then either were treated with saline or ganciclovir (GCV). At 6 weeks post-injury, mice were exposed to random air puffs events while high speed videography recorded whisker and eye movement. MatLab code video processing with publicly available BIOTACT algorithm automatically tracked whiskers.

**Results:** Whisker velocity was calculated using binning statistical analysis. Saline treated animals confirmed our model’s ability to detect aberrant movement such that intact (left) facial nerves caused whisker protraction, while repaired (right) facial nerves had retraction. Administration of GCV increased whisker retraction compared to saline. GCV did not impact intact animal whisker movement compared to repaired whiskers.

**Conclusions:** Inhibition of reactive Schwann cell proliferation appears to worse the degree of synkinesis, providing important insight into a potential therapeutic target for facial nerve injury.

## INTRODUCTION

Facial nerve injury severely limits facial function typically resulting in weak facial expressions and abnormal simultaneous mouth movement and eye closure or synkinesis^1^. This aberrancy can occur either within the nerve fascicles itself or at the facial muscle motor end-plates, resulting in the physiologic finding of synkinesis^2^. Despite advances in microsurgical techniques, complete nerve transection with primary neurorrhaphy is classically believed to lead incomplete recovery with synkinesis.

A well-characterized model to study nerve reinnervation is the mouse femoral nerve transection model^3^. In the femoral nerve model, there are two main branches re-innervating motoneurons can follow, namely the motor branch or the cutaneous (saphenous) branch. To investigate the role of neurotrophic factors on axonal guidance, the transgenic GFAP-thymidine kinase (TK) mouse model was employed. GFAP is expressed solely in reactive Schwann cells and not in uninjured Schwann cell environments^3^ Adminstration of ganciclovir (GCV) ablates these reactive Schwann cells. In the absence of end-organ contact (i.e., tie off distal cutaneous and muscle branches), more motoneurons traveled down the cutaneous branch compared to the motor branch. When examining potential causes for this preference, these authors identified a 2-3-fold increase in reactive Schwann cell proliferation in the cutaneous branch compared to the muscle branch following transection. This suggested that Schwann cell proliferation plays a critical role in directing motoneuron reinnervation. However, they noted, that after GCV application in transgenic GFAP-TK mice (resulting in the removal of immature Schwann cells during reinnervation) this reinnervation preference was eliminated^3^. Moreover, end-organ presence (i.e., patent distal branches) with or without GCV resulted in motoneuron preference to the motor branch, but still some cutaneous projections. These data argue that axonal pathway selectivity is the result of a dynamic interaction between powerful attractant guidance cues emanating from the end-targets as well as reactive Schwann cells lining the distal pathway.

Transgenic mice, GFAP-TK in this case, have demonstrated promising results in femoral nerve regeneration. Here we present our transgenic mouse facial nerve model to study impact of Schwann cell on synkinesis following facial nerve injury and immediate primary neurorrhaphy.

## METHODS

### Nerve transection and drug delivery procedure

This study was approved in accordance to the US Army Institute of Surgical Research (ISR) IACUC protocol # A15-027. Sixteen 6-8 week old (22-27 g) female transgenic GFAP-TK mice were obtained from Jackson Laboratory (stock #005698 Bar Harbor, ME)^4^. Mice were anesthetized with an intraperitoneal injection of ketamine (80-100 mg/kg) and midazolam (4-5 mg/kg). Depth of anesthesia was monitored by a combination of mechanical and observational monitoring techniques including but not limited to: respiratory monitoring, response to stimulus, presence or absence of movement during injury. Fur was removed from the right face and surgical field sterilized. Under microscopic visualization, a post-auricular incision was made and blunt dissection to the main trunk of right facial nerve was identified. Connective tissue was freed from surround nerve. Sharp transection of the nerve at the apex of external auditory canal was made. Fibrin sealant was immediately applied to tension free apposition of nerve ends for anastomosis. The surgical bed was closed with absorbable sutures and the animals were monitored according to ISR post-procedure protocol. Osmotic pumps (Durect Inc, Cupertino, CA) filled either with saline or ganciclovir were placed in a dorsal subcutaneous pocket. Based on nominal performance data from the manufacturer we anticipate that fluid will be delivered at 0.5 μl/hr (± 0.1 μl/hr) allowing for 7 day infusion.Ganciclovir (GCV) was dosed according to Madison et al 20 mg/kg/day diluted in saline. This dose is reported to kill any GFAP-TK expressing cells. At one week animals were sedated to remove osmotic pumps.

### High-speed videography

At six weeks post-operatively, the animals were imaged using high speed videography at 500 frames per second using Fastec TS3 camera (Fastec, San Diego, CA). The camera was positioned directly overhead with lightsource positioned directly underneath A puff of air was presented to the mouse snout positioned 3 cm from a measured and standard puff delivered for 0.1 millisecond using AirStim system (San Diego Instruments, San Diego, CA). Puffs were randomly delivered, to avoid habituation. Using a modified conical tube, the body of the animal was restrained while the head was freely mobile (figure 1). Videography data was analyzed using publicly available MatLab code (Mathworks, Natick, MA) namely the BIOTACT Whisker Tracking Tool (http://bwtt.sourceforge.net). Please refer to this URL to review how whisker position was selected and extracted. Briefly, AVI files were uploaded to the BIOTACT program and whisker position was calculated in radians in each frame. On the right side of screen, animals left face, whisker protraction was represented with positive radian. However, the left side of screen, animal’s right face, whisker protraction was represented with negative radians. Figure 2 illustrates the convention of whisker movement.

**Figure 1:**
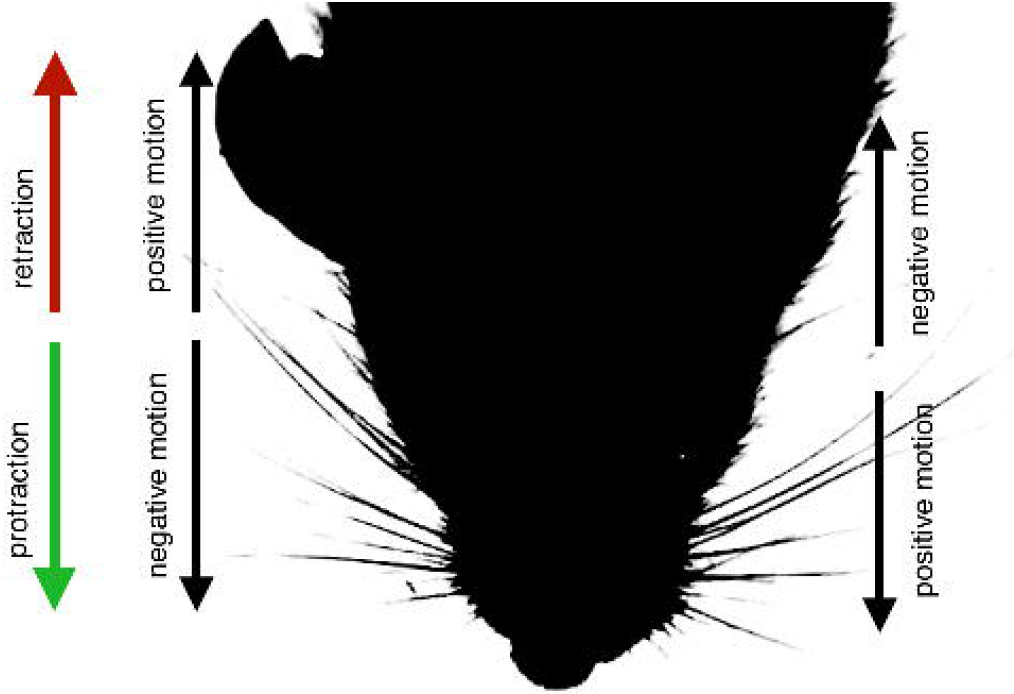
Animal positioning with top-down high-speed videography

**Figure 2:**
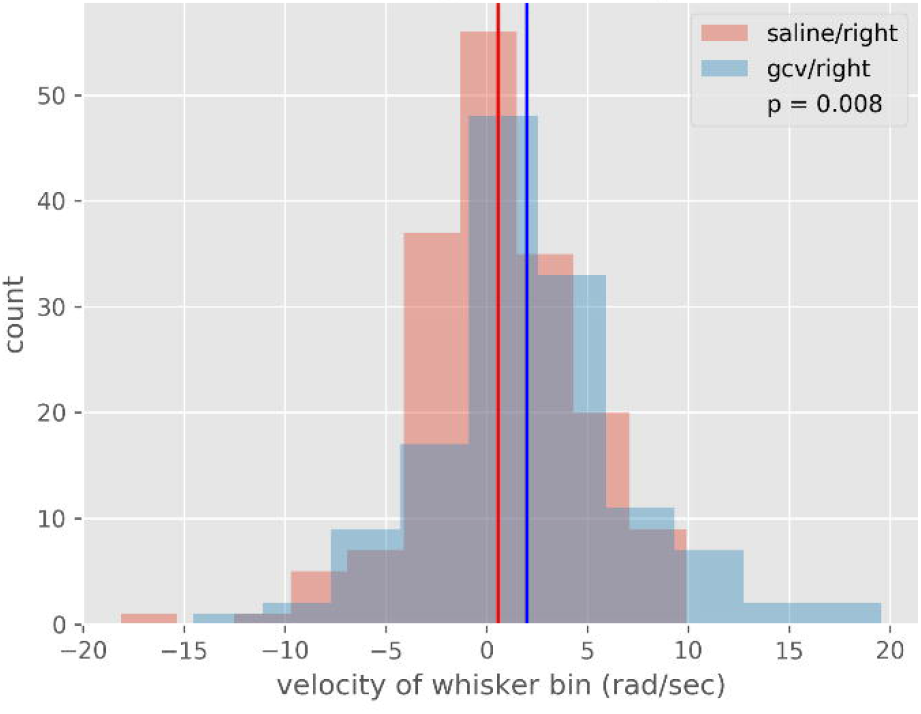
Convention of whisker movement

In the GCV group, a total of 96 separate puffs were analyzed and a total of 111 separate puffs were analyzed in the saline group. Videos were manually annotated to determine the exact moment eye closure began-marking the beginning at which nerve impulse started to close the eye and move the whiskers. This provided the reference point to which whisker movement was determined.

### Statistical analysis

In order to compare the movement of one whisker run to another, we compared the average velocity of the whiskers. This required solving two problems: One, the resolution of the tracking software was not sufficient to track individual whiskers, and adjacent whiskers were often confused. Two, it was not clear how long after the puff the whisker motion was relevant to the question being studied. Too long after, and the motion would be dominated by the stochastic motion which characterizes the whisker motion in an ambient environment, and looking at too few post-puff frames would not give sufficient data to extract reliable statistics.

### The resolution problem

The tracking software we used to determine the whisker’s trajectory would often confuse two adjacent whiskers, especially when the two whiskers overlapped. This meant that the position data was not reliable for any given whisker. However, the tracking software was never in error by more than two adjacent whiskers, due to how they’re grouped on the mouse. This meant that while individual whiskers were unreliable, we could bin the whisker trajectories, and tracking the mean position of a given bin provided a reliable measure of the whiskers contained within that bin.

We found that binning with bins of 5 degrees enabled us to see useful features of the whisker motion (pre- and post-puff stochasticity with a quiescent period immediately post-puff) while also smoothing out the nonphysical tracking features.

### Determining the relevant data

To determine how many frames post-puff we were interested in, we turned to a mathematical technique called Fourier decomposition. What follows is a brief overview of the technique and how we applied it.

Given any time-series signal (in our case, the position of a whisker over time), it is possible to deconstruct that signal into a sum of pure sinusoidal waves different discrete frequencies. By including more and more of these waves, it is possible to recreate a signal to arbitrary precision. This provides a powerful tool: It is of general interest to the physical sciences to be able to say what frequencies are most powerful for a given signal.

By applying a Fourier decomposition to the whisker data, we were able to characterize the motion of the whiskers in each phase of motion. In particular, by looking at which frequencies were the most powerful at any given time, we were able to place a statistical constraint on which frames were most relevant for velocity comparison. In order to determine how many frames the initial post-puff phase lasts, we adopted the following procedure: 1) Establish the peak of the frequency distribution (the Power Spectral Density or PSD) before the puff (PSD_i). 2) Calculate the peak of the PSD for increasing numbers of frames following the puff (PSD_f). 3) Calculate the magnitude of the difference between these two numbers. As the number of frames increases, we expect the PSD_f to diverge from PSD_i until some maximum difference, at which point it will start to return to its previous stochastic motion, and the numbers begin to converge.

When we perform this analysis, we find that this peak occurs at approximately frame 20 across all data runs. Fig. 4b shows an example of this analysis for one data run.

**Figure 3:**
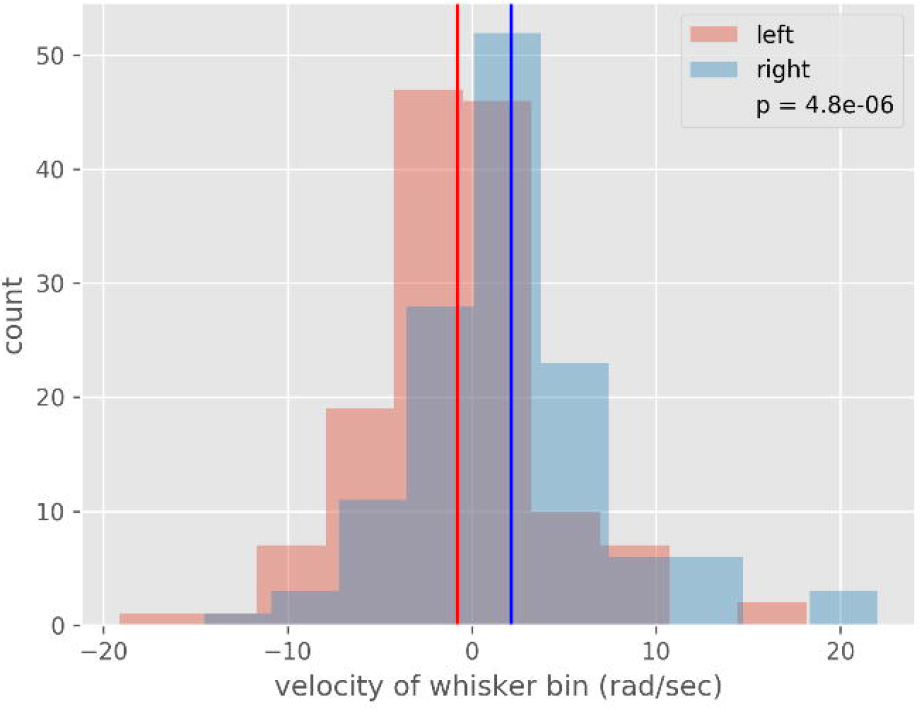
Raw whisker data with puff initiation marked

**Figure 4a:**
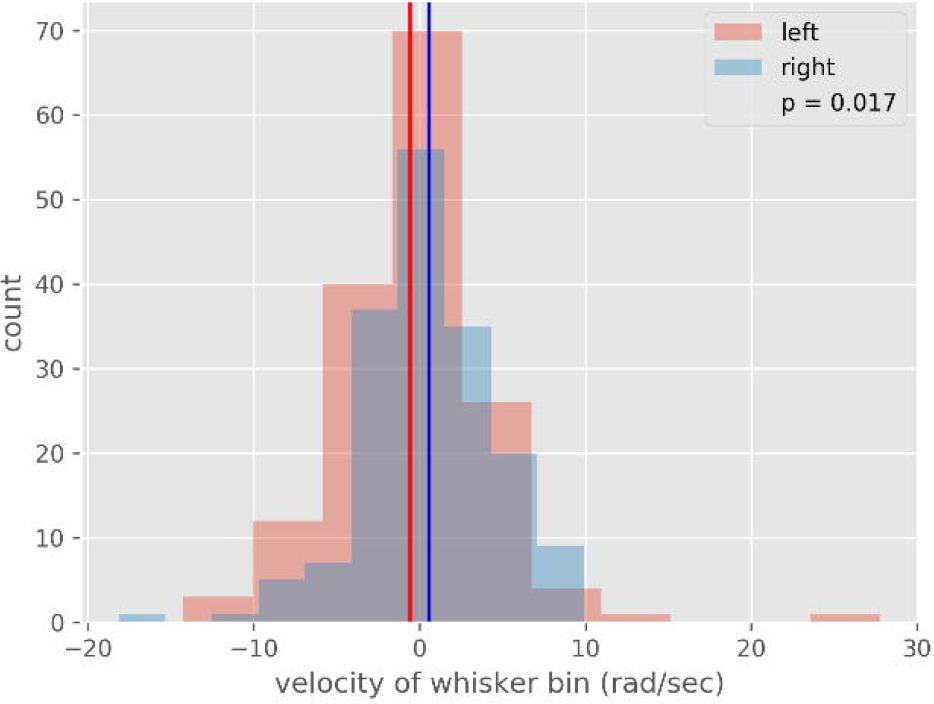
Power spectral density distribution

**Figure 4b:**
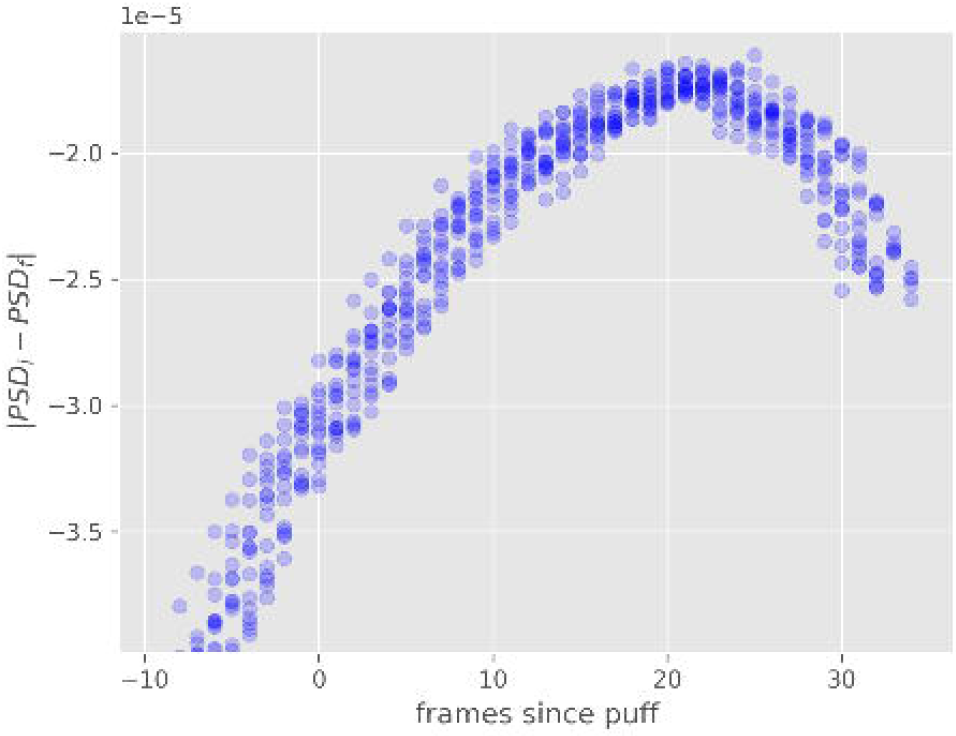
Power spectral density difference demonstrating post-puff peak

Having done this, we could extract the data from just those frames which immediately followed the puff, and look at their average velocity in order to compare the motion of whiskers from different data runs.

## RESULTS

### High speed videography able to capture whisker response following puff stimulus

At six weeks post-injury transgenic mice administered either saline or GCV were recorded with high speed videography to determine if the Biotact algorithm was sensitive enough to capture whisker movement. The raw data was expressed in radians versus frame, with each frame being 1/500 seconds. Figure 3 is sample raw data from saline treated animal.

### Determination of non-stochastic puff response time interval

To evaluate the post-puff velocity behavior, we evaluate the whisker velocity in the 20 frames immediately following the puff. That number was determined by investigating the average number of frames required for the whiskers to return to a pre-puff state. Characterizing the pre-puff motion by looking at the distribution in frequency (known as the power spectral density or PSD), we can compare the post-puff PSD (PSDf) with the pre-puff PSD (PSDi) by making a simple subtraction and taking the average value. As we increase the number of frames since the puff, this difference will grow as the frequencies of the whisker movements experience a randomization period from the puff, until they start to return to their pre-puff distribution. Thus, the difference in PSD pre- and post-puff will show a peak. This is demonstrated for one data run in figure 4a. We use this peak as the threshold to perform the velocity analysis, as this is the data which will best capture the whisker behavior immediately after the puff, before it has returned to its undisturbed (pre-puff) motion. Figure 4b shows the peak for all data runs, showing that the peak occurs at 20.4 ± 3 frames. For simplicity, we take the value to be 20.

### GCV suppression increases synkinetic facial nerve function

Following puff the normal facial nerve response is to simultaneously retract the whiskers and close the eye. Calculating the velocity of whisker movement incorporates speed and direction relative to the start of eye closure. Comparison between the intact and repaired side therefore allows determination of synkinetic movement.

Following puff exposure, the intact side in both saline and GCV administered mice was the same with a mean velocity (p=0.94) (Figure 5a) demonstrating GCV has no impact on intact nerves. In the saline group, velocity response between intact (−0.24 rad/s, protraction) and repaired (0.34 rad/s, retraction) sides was statistically different (p<0.05) (figure 5b). In the GCV group, velocity response between intact and repaired sides was statistically different, such that the right side (mean value of 4.3 rad/s, retraction) is significantly different from the left (mean value −1.2 rad/s, protraction) (p<0.001) (figure 5C). Comparison between saline and GCV groups on the injured side demonstrated mean velocity of 3.2 rad/s (retraction) with GCV treatment versus saline treatment velocity 0.8 rad/s (retraction) (p<0.05) (figure 5D).

**Figure 5a:**
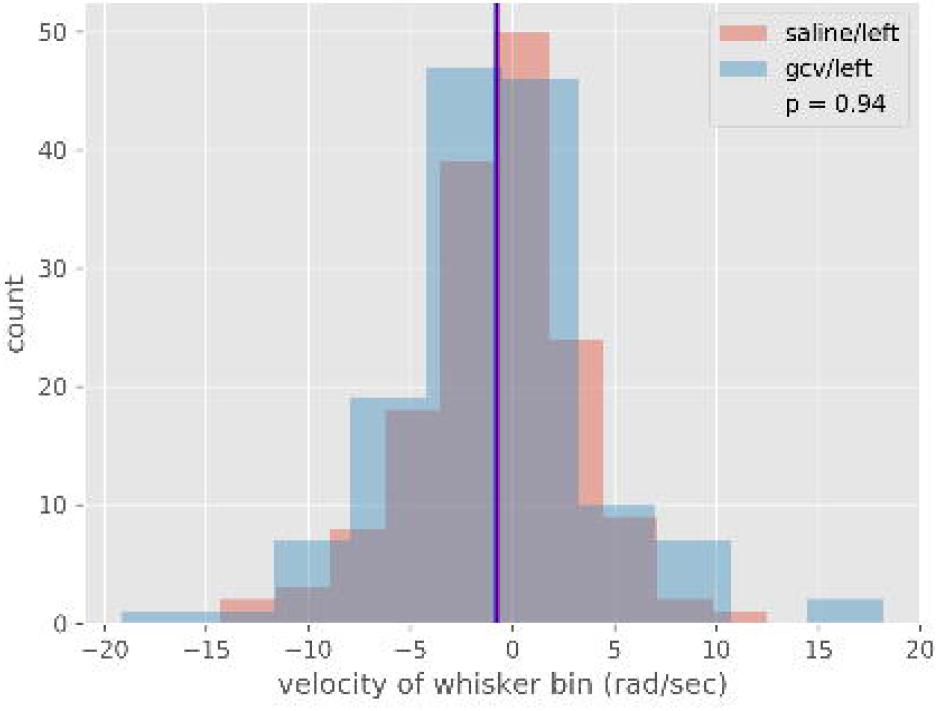
Velocity response on intact side (left) after puff

**Figure 5b:**
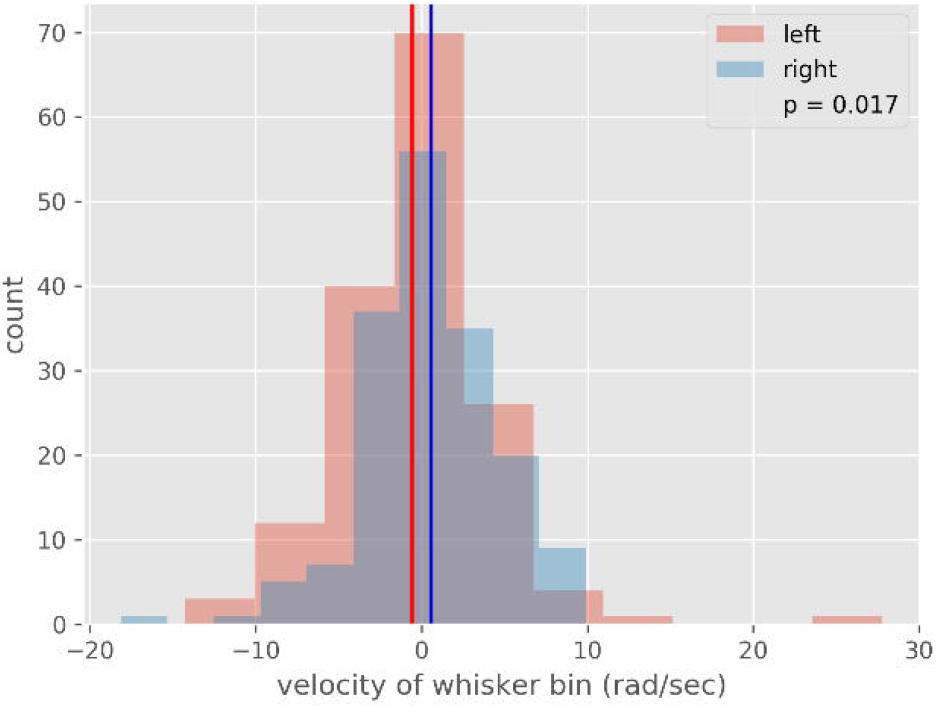
Velocity response with saline treatment

**Figure 5c:**
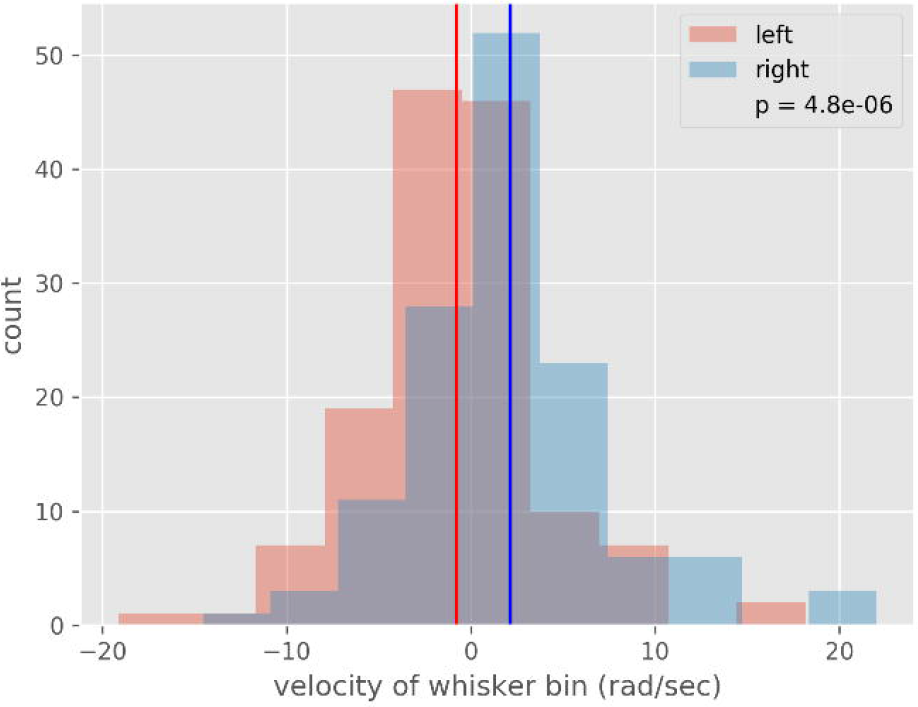
Velocity response with GCV treatment

**Figure 5d:**
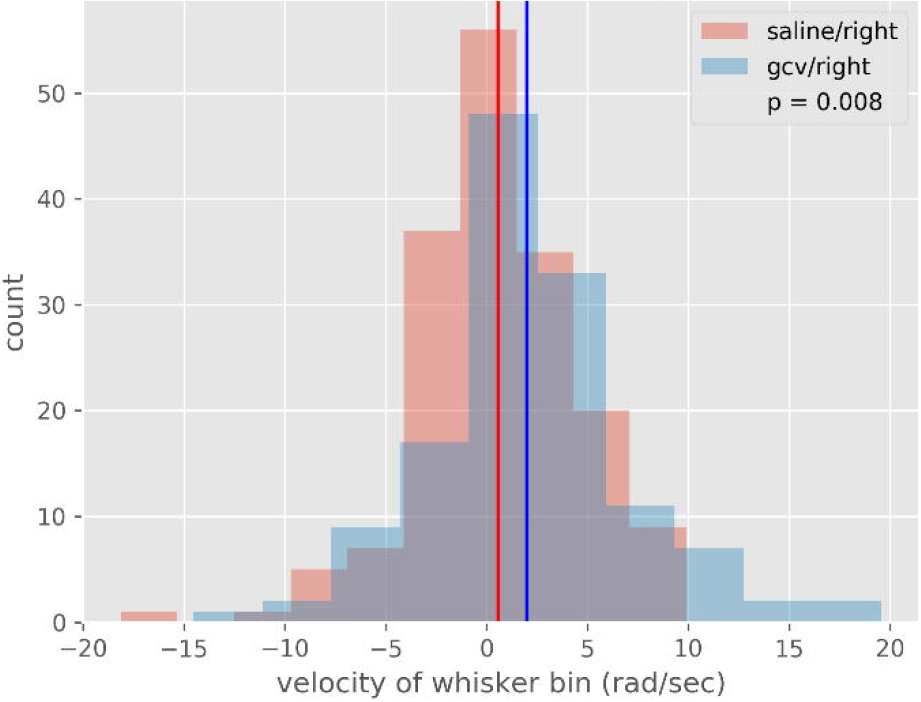
Velocity comparison between saline and GCV treatment on injured side

## DISCUSSION

In this study we administered GCV to mice with the transgenic mutation of GFAP-TK to suppress post-injury Schwann cell proliferation. Our main finding is that reactive Schwann cell proliferation increases synkinetic action. The basis of this study was derived from work by Madison et al where they describe following sciatic nerve injury, adult reactive Schwann cells de-differentiate and re-express glial fibrillary acidic protein (GFAP)^3^.

Defining facial nerve synkinesis is a matter of understanding normal facial nerve function. In the rodent, facial nerve function dictates eye lid and whisker position^5^. Following puff exposure, the intact side responded with eyelid closure and whisker protraction. In the injured side, retraction was seen with eyelid closure representing synkinesis. Suppression of reactive Schwann cells appears to worsen facial nerve recovery, suggesting that they play a key role in regeneration accuracy.

Here we capture facial nerve function with minimal modification to animal itself. Additionally, high spatiotemporal videography (ie 500 frames per second) allowed for large amount of data pertaining to simultaneous eyelid and whisker position. Estimation of error in calculation of whisker position was minimal at 500 frames per second^6^.

The main weakness of our study is the lack of histologic confirmation of GCV suppression of reactive Schwann cell response and success of regeneration through anastomosis and at the muscle end-plates. Additionally, various repair methods other than tissue glue could have been compared. Lastly, tracking temporal change in facial nerve function before and beyond 6 weeks could elucidate progressive worsening or improvement of facial nerve function.

In conclusion, this study develops the first transgenic mouse model to study synkinesis. We demonstrate, the pharmacologic suppression of Schwann negative impact on facial nerve function. The benefit of using a mouse model, is the vast number of genetic mutations that can be targeted for parallel, high through-put post-injury and therapy screening of molecular targets.

